# Primary Studies on the Effect of Microbially Coalbed Methane Enhancement in Suit in the Qinshui Basin

**DOI:** 10.1101/532671

**Authors:** Dong Xiao, Cong Zhang, Enyuan Wang, Hailun He, Yidong Zhang, Junyong Wu, Liuping Deng, Meng Wang

**Affiliations:** State Key Laboratory of Coal Resources and Safe Mining, China University of Mining and Technology, Xuzhou, Jiangsu province, 221116 China; Shanxi Coal Bed Methane Company of Petrochina, Jincheng, Shanxi province, 048000, China; Key Laboratory of Gas and Fire Control for Coal Mines, China University of Mining and Technology, Xuzhou, Jiangsu province, 221116 China; School of life science, Central South University, Changsha, Hunan, 410083 China

**Keywords:** coal bed methane, biogasification, bioenergy, coal, *methanogen sp*., biome

## Abstract

Methods used to yield bio-methane with coal to increase coalbed methane reserves had researched, thus providing a means for improving gas drainage efficiency. One such method utilized to convert coal into gas involves coal biodegradation technology. In order to confirm the practical application of this technology, the experiments were conducted in wells, Z-159, Z-163, Z-167, and Z-7H, in the Qinshui Basin in China, and the duration of the experiments was 32 months. Cl^-^ ion tracer, number changes of *Methanogen sp*., and coal bed biome evolution indicated that the culture medium diffused in the Z-159 and Z-7H wells. These wells resumed gas production separately. Gasification of coal lasted 635 and 799 days, and yielded 74817 m^3^ and 251754 m^3^ coalbed methane in Z-159 and Z-7H wells, respectively. Results demonstrate that coalbed methane enhancement with biogasification of coal is a potential technical to achieve the productivity improvement of coalbed methane wells.

## 1. INTRODUCTION

Coal bed methane (CBM) is a type of associated gas resulting from coal production. The main component of this gas is methane. CBM is a valuable energy resource, meanwhile, it is a dangerous source for mining [1]. Gas drainage represents a widely used technique that not only controls CBM operations but improves the comprehensive utilization efficiency of fossil energy. Therefore, research involving gas drainage mechanisms, applications and coal gasification methods, which service to increase CBM reserves and improve gas well productivity has gradually deepened. [2–4]. Biogasification of coal is one of the methods of coal gasification [5]. The coal components are complex. It contains organic compounds and inorganic components. The methanogenic bacteria could degrade a part of biodegradable organics to convert coal into methane. Activating characteristics of *Methanogen sp*. in coal of the Qinshui basin, and the biodegradability of coal have been preliminary studied in previous research [5]. The in-situ efficiency of this technical requires further research.

The Qinshui basin, located in the southern region of the Shanxi province, has an abundance of CBM and constitutes one of the most important CBM reservoirs and gas development regions in China (Figure. 1). The CBM reserves are estimated at 3.95 trillion m^3^ within this 27137 Km^2^ area [6]. As defined by the Chinese petroleum and natural gas reserves standard, this basin represents a massive gas field. The geologic structure of the Qinshui basin is comprised of the compound syncline, which warps the seams of the southern and northern basins and supplies the fundamental symmetry- bearing portion of the eastern and western regions. The geologic structure in the middle of this basin is flat and contains few faults. The Fm 3#, Fm 9# and Fm 15# coal beds are the main gas bearing layers with good permeability conditions [7].

**Figure 1.**
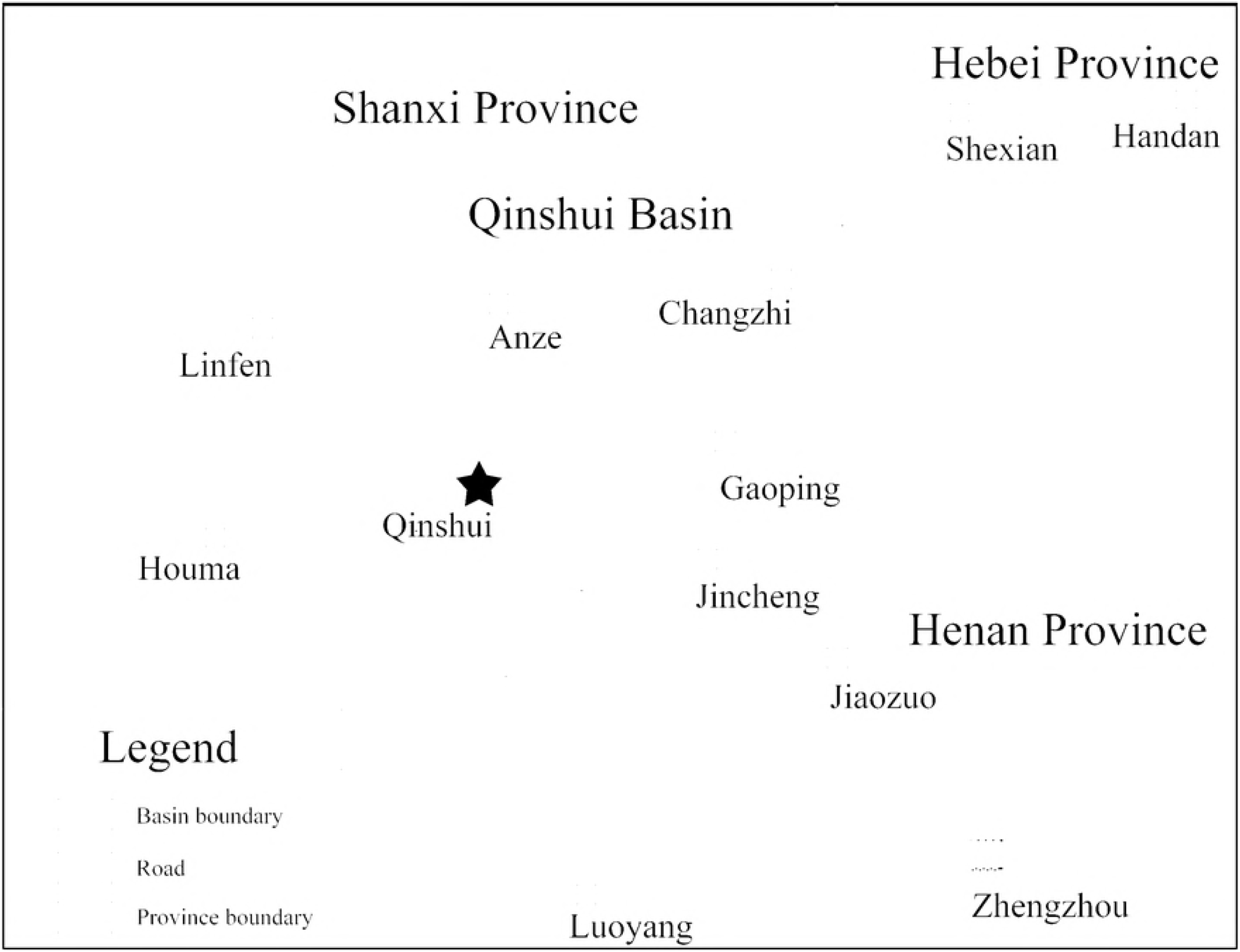
The Qinshui Basin geographical location map. The Qinshui Basin is located in the southern part of Shanxi Province. Neighboring provinces of Herbei and Henan. Anze, Changzhi, Qinshui, Gaoping, and Jincheng are the main coal bed gas development areas. The test location was in Qinshui, where marked with a pentagram symbol in the figure.

Methane productivity of some wells is quite low, and even did not produce gas in a part wells, which like Z-7H, Z-159, and Z-163 wells in experiment. Nitrogen foam fracturing have been used in these wells to expansion crack and improve coal permeability. However, the effect was not satisfying. In this case, the CBM development cost was prohibitive[1]. In previous research, methanogenic bacteria, which include hydrolytic fermentation bacteria, fermenting bacteria, and *Methanogen sp*., etc. had been found in Fm 3#, Fm 9# coal bed in Qinshui basin, and the microbe culture method had been studied also[5,8]. If the coal biogasification technology can be applied on site, it would prove a potential technical to improve gas wells production efficiency and control the cost of CBM development.

## 2. Materials and methods

### 2.1 Experiment Wells and Control Wells Survey

This experiment was carried out in a horizontal multiple branching well and a vertical well. The horizontal well identified as Z-7H (GPS coordinates: 38.716117, 106.475351) and Z-159(GPS coordinates: 35.710277, 112.472500). Z-163, Z-167wells, which were located beside the test wells, were selected as control wells.

Total drill footage of Z-7H well was 5472 m, pure drilling footage in coal was 4405 m and a well control area consisted of 0.371 Km^2^. The connection point depth was 797.5 m, and the deflection point depth was 508 m. The well diameter was 6 inches. The Z-7H well has 2 main well bores and 9 branches. The gently sloping coal seam provides for a favorable geographical condition (Figure. 2). The well was completed on October 15, 2012. The desorption pressure gradually increased and achieved a value of 2.5 MPa and began to product gas on February 23, 2013. However, Z-7H stopped gas production after 644 m^3^ of gas were produced from February to March. The average gas production over the production period was only 64.37 m^3^/day from its initial commissioning until the end of 2014.

**Figure 2.**
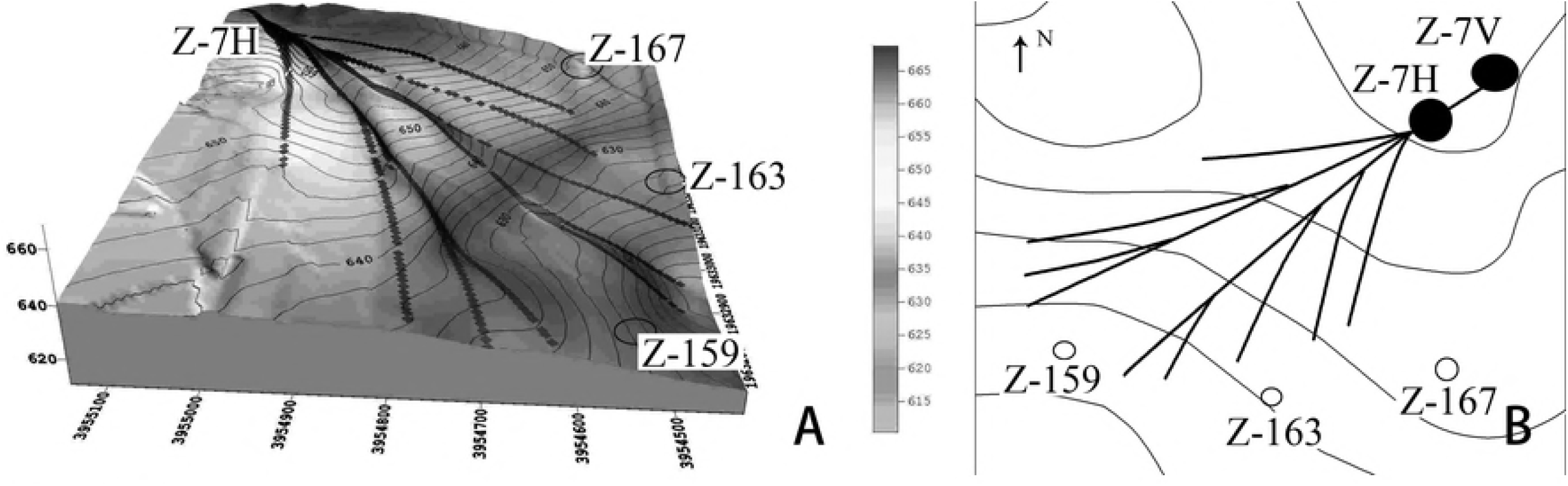
Well Z-7H trajectory stereogram and contour map. Relative geographic locations next to Z-7H well. Wells Z-159, Z-163, and Z-167 are located in south of Z-7H well.

Z-159, Z-163, Z-167 wells were open vertical boreholes, located adjacent to the horizontal pinnate branch well of Z-7H (Figure. 2) and had similar geological conditions to that of Z-7H. The wells diameter were 6 inches, were completed on September to December, 2012, and drilling depth of each borehole were 572.5 m, 591.9 m, 607.9 m respectively. No gas was obtained in Z-159 and Z-163 wells before the experiment performed. Many nitrogen foam fracturing and flushing operations were carried out in Z-7H, Z-159 and Z-163 wells to expansion crack and improve coal permeability, but failed. Well Z-167 had gas productivity of 460 m^3^/day on average before the experiment performed.

### 2.2 Condition of the Fm 3# Coal Seam

Fm 3# coal in Qinshui is a high rank coal with a value of R_o_,_max_>3.0. This coal seam is formed of semibright coal and glance coal, and a small amount of semidull coal. The porosity of FM 3# coal averages 4.88–6.14 %. The main structure in this coal seam is formed by an initial fissure; fissure nondevelopment. The fissure number is 6-11 per 5.0 cm, length is 0.5-10.0 cm, and width is 2.0-120.0 μm. The connectivity of these fissures is low, and is influenced by a calcite film in the coal. The average permeability of Fm 3# coal is 0.02 mD, which was tested using the injection-falloff method in this zone. The reservoir pressure for coal bed gas is 5.0-7.0 MPa, the fluid pressure gradient is 0.07-0.09 MPa/m, and the desorption pressure is 2.0-2.5 MPa. Blockages form easily in this coal seam with a nondevelopment fissure.

### 2.3 Medium Preparation and Injection

A standard medium was used to provide acclimation and culture the *Methanogen sp*. in the coal bed. The final concentrations of the compounds (kg/m^3^) were: yeast extract, 0.50; NaHCO_3_, 0.05; NH_4_Cl, 2.30; KH_2_PO_4_, 1.30; K_2_HPO_4_, 0.70; NaCl, 0.05; MgSO_4_·7H_2_O, 0.20; CaCl_2_·2H_2_O, 0.05. The final pH was 6.80, and the Cl^-^ ion concentration was 44.59 mmol/L [9].

100 m^3^ medium were prepared for Z-7H well, and 30 m^3^ medium was prepared for Z-159 well. The medium prepared with a 50 m3 tank car, and injection performed with a fracturing truck (4150 8X8, Benz, Germany). The injection pressure was adjusted and maintained at less than 4.00 MPa. The Z-159 well and Z-7H well were sealed in March 11 and 26, 2015. And the wellhead pressure was monitored no less than 2 months (Figure 3).

**Figure 3.**
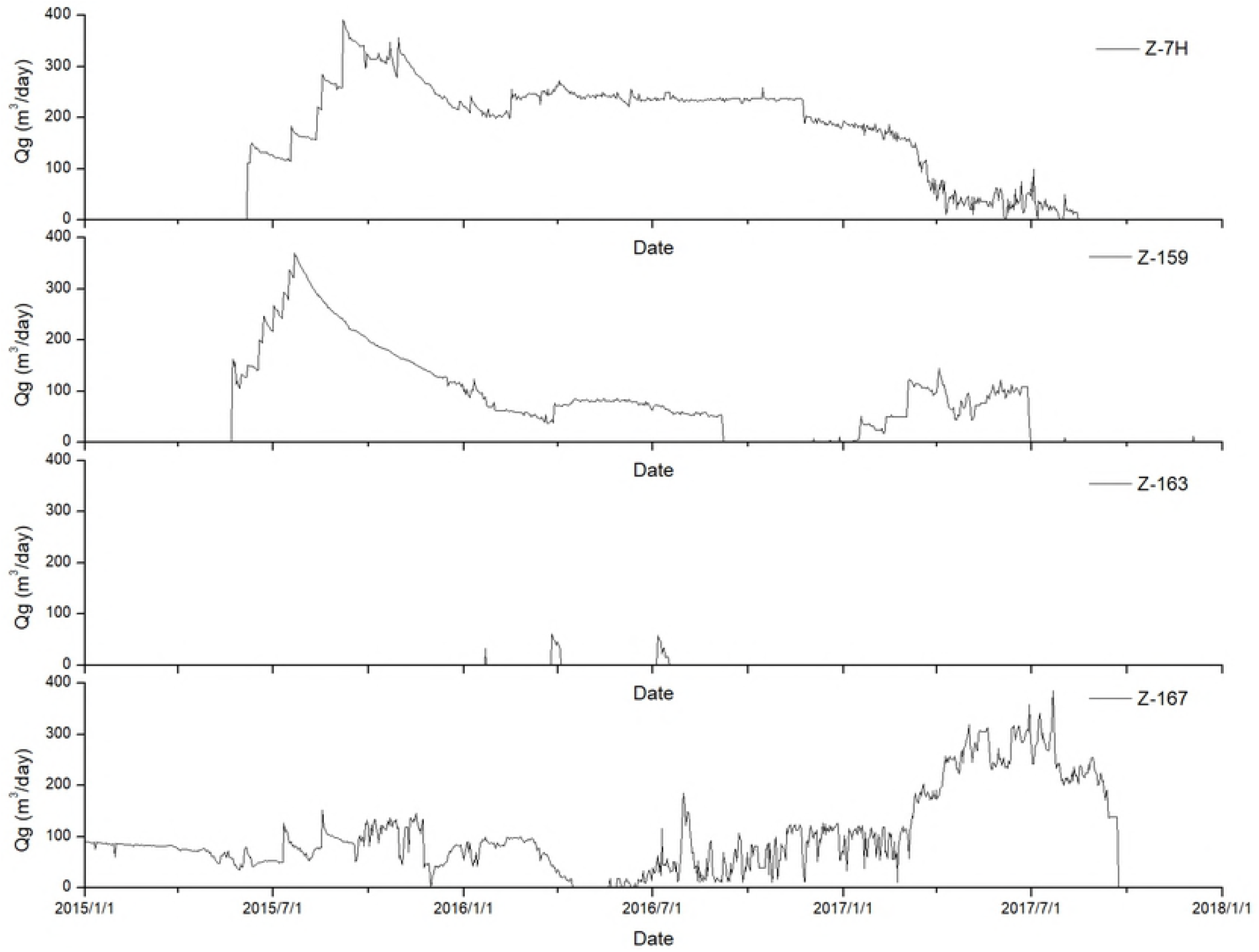
Gas production (Qg) changes in experiment wells and in contrast wells during experiment. The Z-7H and Z-159 were experiment wells. And Z-163 and Z- 167 were contrast wells. The Z-159 well resumed gas production on May 23, 2015 and terminated on February 16, 2017. And the Z-7H wells resumed gas production on June 7, 2015 and terminated on August 14, 2017.

### 2.4 Medium Diffusion Monitors

Medium diffusion in coal seam was monitored by Cl^-^ concentration changes within the underground water [9]. Concentrations of Cl^-^ were identified using an ICS- 1100 ion chromatography system (Thermo Scientific Dionex, Bannockburn, United States). The underground water sample was collected in wells Z-159, Z-163, Z-167 and Z-7H and used aseptic, anaerobic 50 ml tubes. High-speed multifunction centrifugation (J2-MC, Beckman Instruments, Fullerton, United States) and 0.22um filter membrane were utilized to separate the suspended particles and microbes in underground water samples. Purified samples were stored at −80°C in a refrigerator (DW-86L728J, Haier, Qingdao, China).

### 2.5 Microbial observation

Observations of microbe formation, fluorescence detection and microbial counting were performed using an Olympus BX41 (Olympus, Tokyo, Japan) fluorescence microscope at 400× and 1000× magnification with a blood cell counting plate. The F420 fluorescence method was used to test for the presence of methanogenic bacteria (McInerney and Beaty, 1988,).

### 2.6 CBM wells productivity monitor

CBM production; methane, carbon dioxide, and nitrogen concentrations were the parameters used to analyze gas productivity and identify the effects of biological coal gasification. Gas production rates in each well were monitored using a gas roots flow meter (model: FLLQ, Fuma, Wenzhou, China). Measurement accuracy of the FLLQ gas roots flow meter was class 1.0, and flow range of this meter was 0.6-400 m3/h.

Methane and carbon dioxide analyses were performced using Agilent 7890A gas chromatograph (Agilent, Tokyo, Japan). The nitrogen (carrier gas) flow rate was fixed at 1 ml/min. The injection port was maintained at 150 °C, the oven temperature was 25 °C and the TCD was operated at 200 °C. Retention time for methane was 3.76 minutes and 5.0 minutes for carbon dioxide. Calibration standards consisting of 40% methane, 20% carbon dioxide, 10% hydrogen and 30% nitrogen were injected at atmospheric pressure to provide the calibration plot.

### 2.7 DNA extraction and PCR

100 mL of underground water in the Z-7H well was collected at the initial and termination stages of this experiment. Bacteria was concentrated to 1 mL by centrifugation (J2-MC, Beckman Instruments, Fullerton, United States) and stored in cryovials at −80°C (refrigerator type: DW-86L728J, Haier, Qingdao, China) until DNA was extracted. The centrifugal force was set to 13000 X g, and centrifuged for 10 minutes. Total genomic DNA was extracted from 1 mL concentrated underground water samples using E.A.N.A. Soil DNA Kit (OMEGA, Georgia, GA, USA) following the manufacturer’s instructions.

The V4 region of 16S rRNA gene was amplified with polymerase chain reaction (PCR) using primers 515F (5’- GTG CCA GCM GCC GCG GTAA - 3’) and 806R (5’- GGA CTA CHV GGG TWT CTA AT - 3’) [11]. The primer pair was reported to generate an optimal community clustering with the sequence length in the V4 region [12]. Each 20 μL PCR reaction was composed of 2 ng of template DNA, 0.2 μM primers, 0.2 mM dNTP, 2 μL 10×Pfu Buffer with MgSO_4_ (Applied Thermo, Califonia, CA, USA), with H_2_O up to 20 μL. The DNA amplification was performed under the following cycling conditions: 1 cycle of 2 min at 95 °C, followed by 30 cycles with 30 s at 95 °C, 30 s at 55 °C and 1 min at 72 °C, then a final extension period of 5 min at 72 °C. Before sequencing on the Illumina Miseq sequencing platform, we would amplify the V4 region by adding sample-specific 10-base barcodes and universal sequencing tags by Sample-Specific PCR protocol. The PCR procedure was as following: 1 cycle of 95 °C at 2 min, 15 cycles of 95 °C at 15 s, 60 °C at 30 s, 68 °C at 1 min, the final step was 1 cycle of 68 °C at 3 min. Equal volume of each barcoded product was pooled into amplicon libraries and purified using Agencourt AMPure XP system (Beckman Coulter, CA, USA) then examined on Agilent Bioanalyzer 2100 for product size distribution. The purified libraries were quantified with Qubit^®^ dsDNA HS Assay Kit (Life Technologies, CA, USA) and used for sequencing.

### 2.8 Sequencing and Data Analysis

16S rRNA gene libraries were sequenced using an Illumina MiSeq (San Diego, CA, USA) platform and the sequencing data were base-called and demultiplexed using MiSeq Reporter v.1.8.1 (Illumina, SanDiego, CA, USA) with default parameters. The adapter sequences and low quality reads were trimmed away from the raw reads with Trimmomatic v.0.32 [13]. Then the clean reads were analyzed using the Uparse [14] and Mothur pipeline [15] to generate operational taxonomic units (OTUs). The OTUs were picked out at 97% similarity [16]. The resulting representative sequence set was aligned against the core sequence database of the SILVA 123 release with the Mothur script (www.mothur.org/) and given a taxonomic classification using RDP at the 80% confidence level [17]. The richness estimators (ACE and Chao1) and diversity indices (Shannon and Simpson) were calculated using the Mothur program. OTUs’ comparisons were performed using the Venn diagram package. Neighbor-joining phylogenetic tree was used to investigate the similarity of species abundance using the Unweighted Pair Group Method with Arithmetic mean (UPGMA) clustering method [18].

## 3. Results and Discussion

### 3.1 Medium Diffusion in Coal Bed

The key factor needed to enhance the biodegradation of coal is nutrient distribution in coal. Z-159 and Z-7H were the medium injection wells in the experiments. And the medium was injected on March 10 and 25, 2015 respectively. Meanwhile, the Z-163 and Z-167 wells were the contrast wells. The coal bed water samples were collected before the medium was injected, and on 7 February, 2016, after the Z-159 and Z-7H wells resumed gas production. The change in of Cl^-^ ion concentration was served as an indicator to assess whether the medium had diffused in surrounding wells field.

Chloride, in the form of the Cl^-^ ion, constitutes one of the major inorganic anions in the underground water [19,20]. It originates from the dissociation of salts, such as sodium chloride or calcium chloride, in underground water [21]. These salts, and their resulting chloride ions, come from coal seam minerals. Medium diffusion in coal changed the Cl^-^ ion concentration in the underground water, and as a result of anthropogenic impacts [22]. The concentration of Cl^-^ in the underground water of wells Z-7H and Z-159 was altered following nutrition medium injection, showing significant variations in Cl^-^ concentrations due to the effects of nutrition distribution. Cl^-^ concentrations would increase in other wells if there was nutrition seepage flow into the control area of neighboring wells.

Baseline Cl^-^ concentrations for the Z-159, Z-163, Z-167, and Z-7H wells were considered to be stable at 100~110 mg/L. This value increased two to three times in 70 days, after the experiments were performed for the Z-159 and Z-7H wells. In contrast, the Cl^-^ concentration within the Z-163 and Z-167 wells was maintained at about the same level over the experimental period (shown in Table 1). These findings confirmed that the distribution of the culture medium and the coal biogasification were active in the Z-159 and Z-7H wells, and did not spread to the Z-163 and Z-167 wells.

**Table 1.**
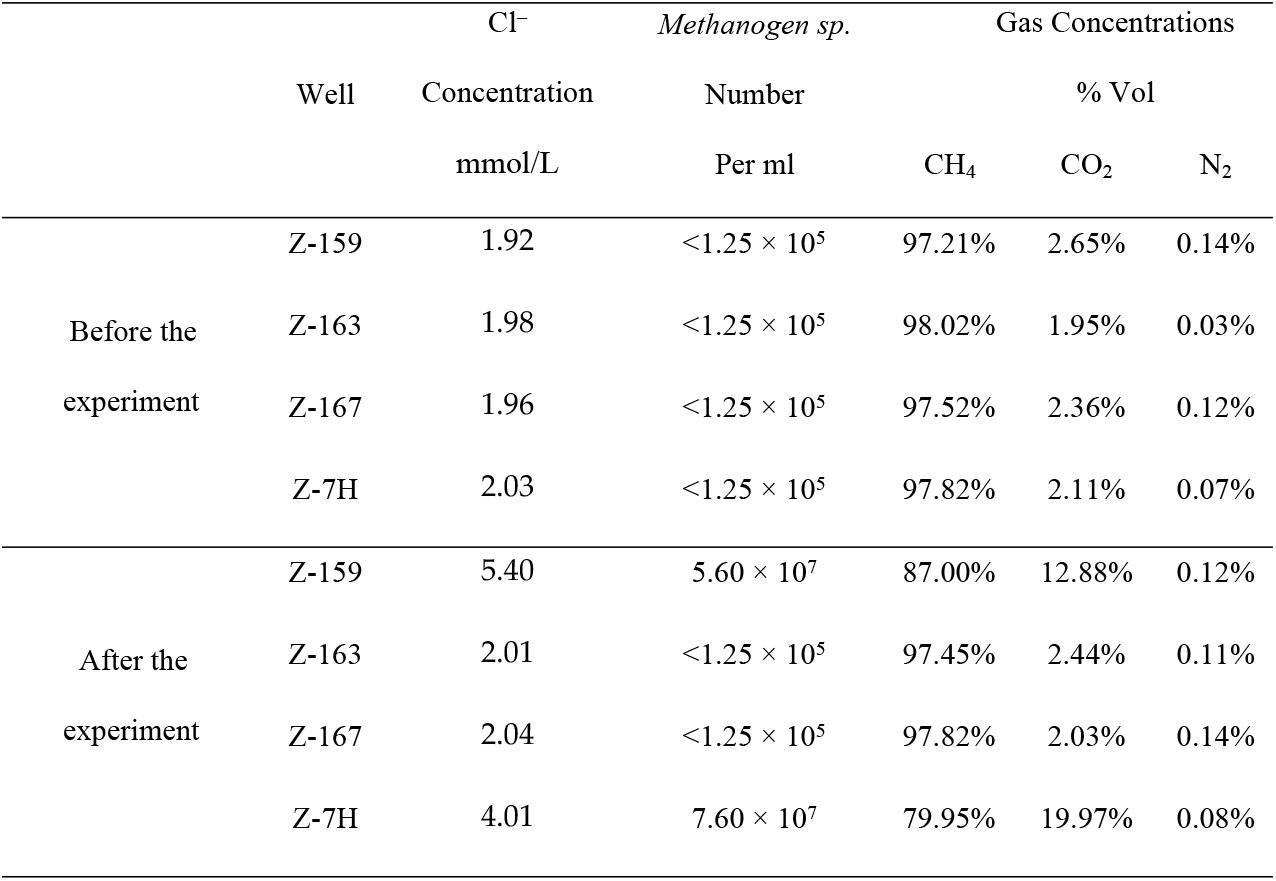
Cl^-^ concentration, *Methanogen sp*. number, and gas concentrations changes, before and during the experiments.

### 3.2 Changes in the number of *Methanogen sp*

The biogasification of coal is performed through the collaboration of different microorganisms. They include hydrolytic fermentation bacteria, fermenting bacteria, and *Methanogen sp*., etc [19]. And these bacteria had been found in Fm 3#, Fm 9# coal bed in Qinshui basin [5,8]. As the terminal bacteria of coal biogasification process, *Methanogen sp*. can indicate the development of methanogenic consortia for coal biodegradation in coal bed [10]. The number of *Methanogen sp*. in the sample could be observed by fluorescence microscopy, because of *Methanogen sp*. has a fluorescent effect in 420 nm light[5,20]. On this condition, it has a positive correction between biogasification of coal with the number of *Methanogen sp*.. Quantitative analyses of *Methanogen sp*. were performed before the medium was injected, and on 7 February, 2016 when the Z-159 and Z-7H wells resumed gas production. Initial levels of *Methanogen sp*. were lower than 1.25 × 10^5^ per ml on average in the experimental and contrast wells. The *Methanogen sp*. numbers increased from less than 1.25×10^5^ to 5.60×10^7^ and 7.60×10^7^ per ml in the Z-159 and Z-7H wells with the medium diffusing in the coal bed (shown in Table 1). Meanwhile, the number was stable at the original level in the Z-163 and Z-167 wells. These findings confirmed that the methanogenic consortia developed with intervention, and created biological conditions for biogasification of coal.

### 3.3 Effect Analysis of biogasification of coal

Biogasification of coal was performed through the bio-fermentation of organic compounds in the coal. 100 m3 and 30 m3 of medium were injected into the Z-7H and Z-159 wells, and were used to provide main nutrients for microbial growth in the coal. The Z-159 well resumed gas production on May 23, 2015 and terminated on February 16, 2017. And the Z-7H wells resumed gas production on June 7, 2015 and terminated on August 14, 2017. The gasification of coal lasted 635 and 799 days, and yielded 74817 m^3^ and 251754 m^3^ CBM in Z-159 and Z-7H wells, respectively. The gas production of contrast wells maintained the original characteristics during the experiment (Figure 3). Especially, the Z-163 well maintained at non-gas productive state during gas productivity has resumed in experimental wells.

Ion tracers indicate that the nutrition diffused in the Z-159 and Z-7H wells in experiments. With the effect of nutrition control, *Methanogen sp*. numbers increased, from lower than 1.25 × 10^5^ to more than 5.60 × 10^7^ per ml, in nutrition injected wells. These data were found to have a positive correlation with the changes in gas productivity in certain wells.

In this experiment, gas concentrations showed certain changes, which were influenced by the bio fermentation of coal. The yield of carbon dioxide, which as by products of coal biogasification, would influenced the CBM composition. Therefore, the increase in the CO2 content of the CBM could be another indication of the biogasification phenomenon of the coal (indicated in Table 1). The gas components of CBM in Z-167, which without influence of medium injection, were maintained at a stable level.

### 3.4 Analyses of changes in microbial community structure

Different microbial populations and coal constitute a coal bed ecological community. Microbial community ecology evolution is closely related to changes of coal bed environment factors. These factors include gas composition, nutrition, and specific surface area of coal [8]. Biodiversity testing before and after the experiment in Z-7H well had confirmed this evolution. Data indicated that microbial species have significant changes with medium injection in coal.

**Figure 4.**
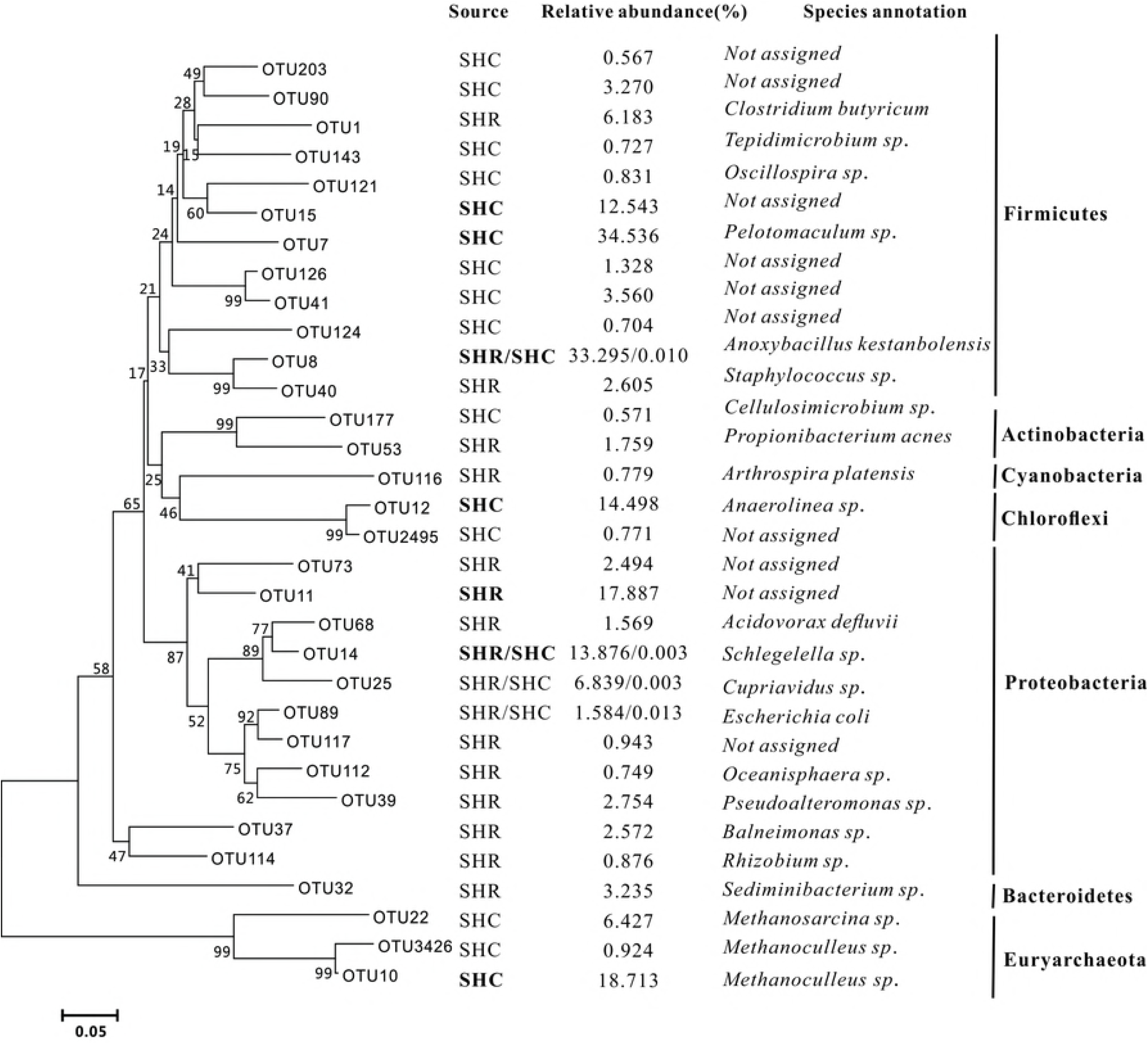
Coalbed microbial community structures before and after experiment. SHR is microbial in row coal bed before experiment. And SHC is coal bed microbial after cultured.

There were 105 strains detected in high-throughput sequencing of the original coal bed microbial community before the experiment. High relatively abundant genus was as follows: *Propionibacterium sp., Balneimonas sp., Anoxybacillus sp., Cupriavidus sp., Schlegelella sp., Clostridium sp., Clostridium sp., Pseudoalteromonas sp., Pseudoalteromonas sp*. etc. After cultured, meanwhile, the detected strains value in High-throughput sequencing of the Z-7H well biome decreased to 85. And compare the microbial community structure before and after experiment, only 11 species were the common specie. And the number of common species with a relative abundance higher than 0.5% is only four, which include: *Anoxybacillus_kestanbolensis sp., Escherichia_coli sp., Cupriavidus sp., Schlegelella sp*.. Analysis of the response of coal bed biome to environmental changes and community evolution showed that two groups of microorganisms were the most obvious. First of all was *Methanoculleus sp*. and *Methanosarcina sp*., both of them belong to the family *Methanomicrobia sp*.. Most *methanogen sp*. make methane from CO_2_ and H_2_. Others utilize acetate in the acetoclastic pathway. In addition to these two pathways, species of *Methanosarcina sp*. can also metabolize methylated one-carbon compounds through methylotrophic methanogenesis. Such one-carbon compounds include methylamines, methanol, and methyl thiols [21]. Only *Methanosarcina sp*. species possess all three known pathways for methanogenesis, and are capable of utilizing no less than nine methanogenic substrates, including acetate [22]. *Methanoculleus sp*. is differ from other methanogens, it can use ethanol and some secondary alcohols as electron donors as they produce methane [23]. And followed by *Anaerolinea sp*. and some species from *Clostridiales sp.. Anaerolinea sp*. has a fermentative metabolism, utilizing carbohydrates as well as proteinaceous carbon sources [24]. It produces acetate, lactate and hydrogen as by products of glucose fermentation [25]. For *Clostridiales sp., Peptococcaceae sp*. and *Clostridiales sp*., they could produce butyric acid with organic fermentation. Varying concentrations of acetic acid, lactic acid and/or ethanol, propanol or butanol are also formed as fermentation products [26]. They provided nutrients for methanogens and played an important roles in the new biome. The relative abundance of these microbe was very low in original coal bed microbial community. The evolution of this coal bed ecological community demonstrates the critical role of this experimental methods in improving the gas production capacity in situ.

## 4. Discussion

Results from this research have demonstrated that biogasification of coal could be utilized to enhance gas well productivity. The factor of Cl^-^ concentration changes had been tested to assess the medium diffusion in coal. The baseline of this factor was stable at 100-110 mg/L in original coal seam. This value increased two to three times after experiment performed for 70 days in both experimental wells. This conclusion confirmed that the injected medium was maintained within the experimental wells and not diffused to the periphery. Furthermore, the range of nutrition influence was controlled in the scope of Z-159 and Z-7H wells.

Quantitative analyses of *Methanogen sp*. were performed before and after experiment with 420 nm fluorescence counting. The *Methanogen sp*. numbers increased from less than 1.25×10^5^ to 5.60×10^7^ and 7.60×10^7^ per ml in the Z-159 and Z- 7H wells with medium diffusing in the coal bed. The number changes of *methanogen sp*. confirmed that the methanogenic consortia developed with intervention, and created biological conditions for biogasification of coal. The resumption of gas production in wells Z-159 and Z-7H confirmed the effect of CBM enhancement with biogasification of coal. The gasification of coal lasted 635 and 799 days, and yielded 74817 m^3^ and 251754 m^3^ CBM in Z-159 and Z-7H wells, respectively.

High-throughput sequencing had been used to analysis the coalbed ecological community evolution in experiment. The proliferation of *Methanomicrobia sp., Anaerolinea sp. and Clostridiales sp*. reflected the coalbed microbial community changes with medium diffusion in coal. And the biome indicated that the methanogenesis mechanism of the Sihe coalbed had a composite character.

## 5. Acknowledgements

The authors’ acknowledge the contributions of the following companies for allowing access to coal samples and other information used in this paper: Huabei Oil Field, Fenghuangshan mining, J&D Technology Company. We thank Dr. Aikuan Wang and Jiang He, China University of Mining and Technology, for instructions on data analyses. This work was supported by the State Key Laboratory of Coal Resources and Safe Mining Subject (grant number SKLCRSM17KFA08 and SKLCRSM19X012), the Fundamental Research Funds for Central Universities (grant number 2014QNB41).

## Abbreviations

CBM: Coal bed methane;
TCD: Thermal Conductivity Detector.

